# *Lgr5*+ intestinal stem cells are required for organoid survival after genotoxic injury

**DOI:** 10.1101/2024.04.08.588400

**Authors:** Joseph Lee, Antoine Gleizes, Felipe Takaesu, Sarah F Webster, Taylor Hailstock, Nick Barker, Adam D Gracz

**Affiliations:** Department of Medicine, Division of Digestive Diseases, Emory University; Graduate Program in Biochemistry, Cell and Developmental Biology, Emory University; Institute of Molecular and Cell Biology, Agency for Science, Technology and Research (A*STAR), Singapore; Department of Physiology, Yong Loo Lin School of Medicine, National University of Singapore

## Abstract

Progenitors and mature cells can maintain the intestinal epithelium by dedifferentiation and facultative intestinal stem cell (fISC) function when active ISCs (aISCs) are lost to damage. Here, we sought to model fISC activation in intestinal organoids with doxorubicin (DXR), a chemotherapeutic known to ablate *Lgr5*+ aISCs *in vivo*. We identified low and high doses of DXR compatible with long-term organoid survival. Similar fISC gene activation was observed between organoids treated with low vs high DXR, despite significantly decreased survival at the higher dose. aISCs exhibit dose-dependent loss after DXR but survive at doses compatible with organoid survival. We ablated residual aISCs after DXR using a *Lgr5*^*2A-DTR*^ allele and observed that aISC survival of the initial genotoxic insult is required for organoid survival following DXR. These results suggest that while typical fISC genes are activated by DXR injury in organoids, functional stemness remains dependent on the aISC pool. Our data establish a reproducible model of DXR injury in intestinal organoids and reveal differences in *in vitro* responses to an established *in vivo* damage modality.

## INTRODUCTION

The intestinal epithelium is maintained by a pool of *Lgr5*^*high*^ active intestinal stem cells (aISCs), which replenish all mature intestinal epithelial cells (IEC) lineages during homeostasis and are located at the crypt base (Barker et al., 2007). Early observations that the rapidly proliferating cells of the intestinal crypts are sensitive to genotoxic radiation injury motivated research into the existence of “reserve” or “quiescent” ISC populations, which could compensate for loss of the aISC pool. More recently, a consensus emerged that “reserve” ISC function is maintained by facultative ISCs (fISCs), consisting of multiple partially differentiated or fully mature IEC lineages capable of reverting to an ISC state in the setting of injury and regeneration (Meyer et al., 2022). fISCs are characterized by high levels of YAP signaling and the expression of several transient, injury-induced biomarkers, including Sca-1/*Ly6a* and *Clu* (Ayyaz et al., 2019; Nusse et al., 2018). While the induction of fISCs by injury is well-studied, much less is understood about how the epithelium returns to homeostasis and whether long-term impacts of injury remain following regeneration.

Primary intestinal epithelial organoids have emerged as a valuable *in vitro* tool for understanding ISC biology. Organoids are especially promising in context of personalized medicine, where screens using patient-specific organoids may predict responses to therapy and facilitate *in vitro* comparisons of normal and diseased epithelium (Takahashi, 2019). Here, we set out to define a reproducible protocol to model genotoxic aISC injury in intestinal organoids with doxorubicin (DXR), an anthracycline used to treat a wide range of primary and metastatic tumors (Minotti et al., 2004). DXR targets proliferating cells by inhibiting topoisomerase II and evicting histones (Pang et al., 2013). In the intestine, this primarily affects aISCs and results in fISC-driven regeneration (Jones et al., 2019; Sheahan et al., 2021). We reasoned that an optimized DXR injury assay in organoids could model injury-induced cycles of aISC/fISC fate transitions in an epithelial autonomous manner. We found that organoids depend on *Lgr5*+ aISCs for post-DXR survival, raising important considerations for interpretating ISC and regeneration data from organoid assays.

## RESULTS & DISCUSSION

### Reproducible modeling of IEC injury and recovery with optimized DXR treatment

We first sought to identify doses of DXR treatment that could induce expected aISC injury phenotypes while allowing long-term organoid survival, facilitating studies of epithelial recovery. Jejunal organoids from adult mice were allowed to establish in ENR media for 48hr prior to treatment with varying doses of DXR for 4hrs, after which media was changed to prevent ongoing DXR exposure (Fig S1A). We found that 0.25μg/mL DXR was required to induce a significant decrease in organoid survival, while organoids were unable to tolerate treatment with 1.00μg/mL or more, as indicated by only rare surviving organoids at 6d post-DXR (Fig S1B & C). Survival began dropping off at 24hr post-DXR but continued to 5d post-DXR at all tested doses. This suggested a secondary effect of injury, such as loss of ISC function, as opposed to non-specific cell death.

Next, we tested the reproducibility of 0.25μg/mL and 0.50μg/mL DXR injury by quantifying post-DXR survival in 4 independent biological replicates. We made minor modifications to our DXR treatment protocol to facilitate reproducible dosing, including careful mixing of diluted DXR into sample wells and multiple post-DXR rinses (see *Methods*). DXR treatment consistently resulted in loss of budding and shedding of dead cells into the luminal space and surrounding the organoids, with 0.50μg/mL DXR inducing more severe morphological phenotypes (Fig 1A). Variability in post-DXR survival between organoid preps from different mice was minimal (Fig 1B). Additionally, we quantified long-term post-DXR organoid viability and saw no further reduction in survival between 5 and 10 days post-DXR (Fig 1B). These data suggest that the initial “pulse” of 0.25 μg/mL or 0.50μg/mL DXR induces dose-dependent injury that is compatible with long-term organoid survival.

**Figure 1:**
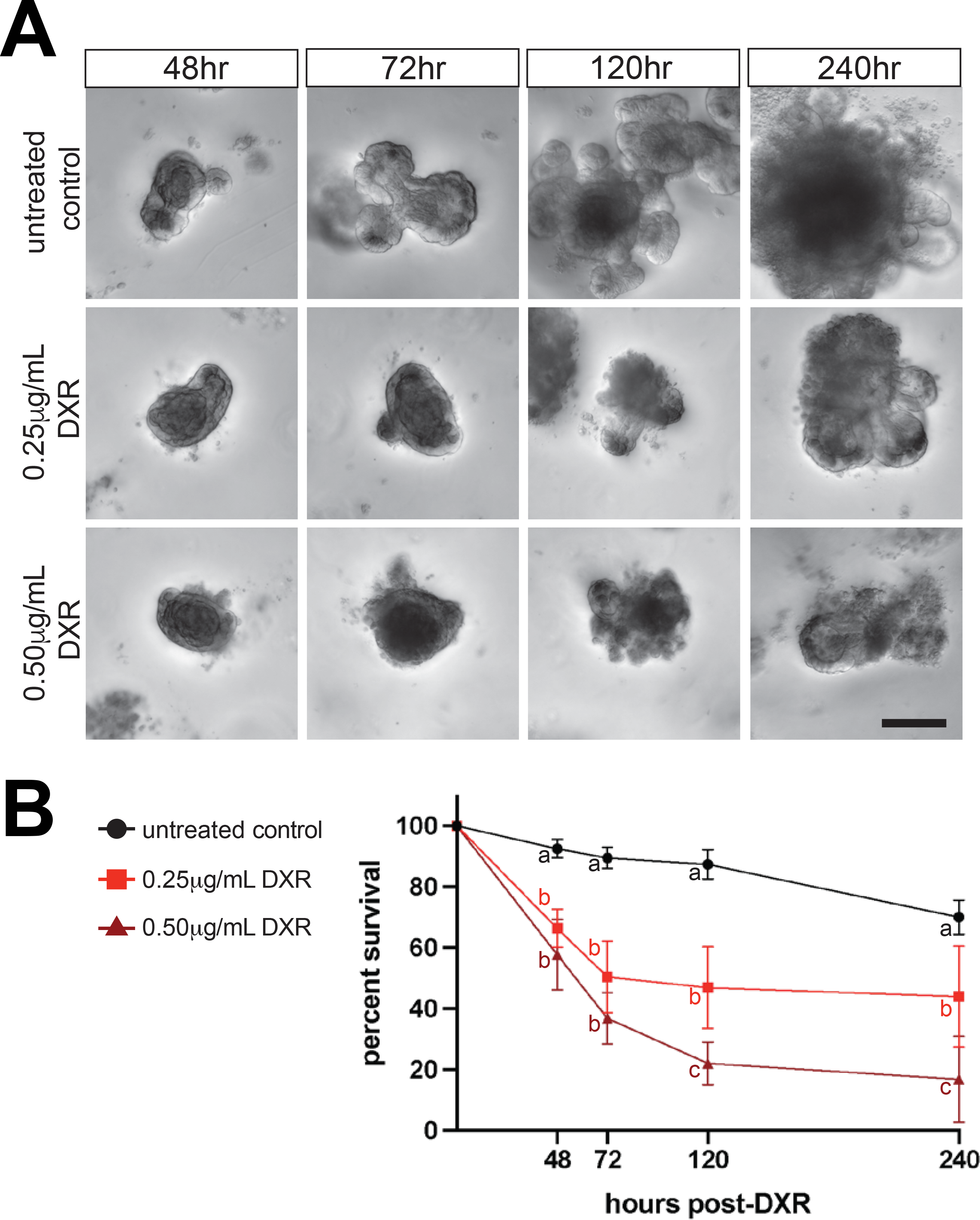
Doxorubicin induces reproducible organoid injury compatible with long-term survival. (A) Organoids treated with 0.25μg/mL and 0.50μg/mL DXR exhibit dose-dependent morphological changes consistent with IEC apoptosis and injury (scale bar represents 100μm). (B) DXR reproducibly decreases organoid survival in a dose-dependent manner and allows some organoids to survive for at least 10d post-injury (n = 3 replicates each from 4 independent organoid preps, different letters indicate significant differences between sample groups at each timepoint, p < 0.05).

### Doxorubicin treatment induces fISC genes and suppresses aISC markers

*In vivo*, aISCs are well-established as the target of DXR injury. Reports demonstrate loss of (1) OLFM4+ CBCs and (2) lineage-tracing of *Lgr5*+ aISCs, following a single dose of DXR (Dekaney et al., 2009; Sheahan et al., 2021). To confirm loss of aISCs and activation of fISC genes, we isolated RNA from untreated controls and organoids treated with 0.25μg/mL and 0.50μg/mL DXR at 24hr and 48hr. Consistent with *in vivo* reports, we observed a significant decrease in expression of *Lgr5* and *Olfm4* at both timepoints, suggesting a rapid loss of aISCs that does not recover by 48hr post-injury (Fig 2A). fISC markers *Ly6a* and *Clu* were significantly upregulated in both doses at 24hr post-DXR (Fig 2B). Interestingly, both fISC markers continued to increase between 24hr and 48hr following 0.50μg/mL DXR, while they did not after 0.25μg/mL DXR. *Ascl2*, which is expressed in aISCs but also mechanistically required for intestinal regeneration in fISCs, was downregulated at 24hr post-DXR, but less affected than other aISC markers (Fig S2) (Murata et al., 2020). Because *Ascl2* decreased between 24hr and 48hr in control organoids, there was no significant difference between controls and DXR-treated organoids at 48hr (Fig S2). These data demonstrate that DXR induces expected loss of aISC genes and induction of fISC markers. Further, higher dose DXR induces a greater upregulation of fISC markers. Together, gene expression at early timepoints suggest that intestinal organoids model the expected aISC/fISC dynamics in response to a single, 4hr pulse of DXR.

**Figure 2:**
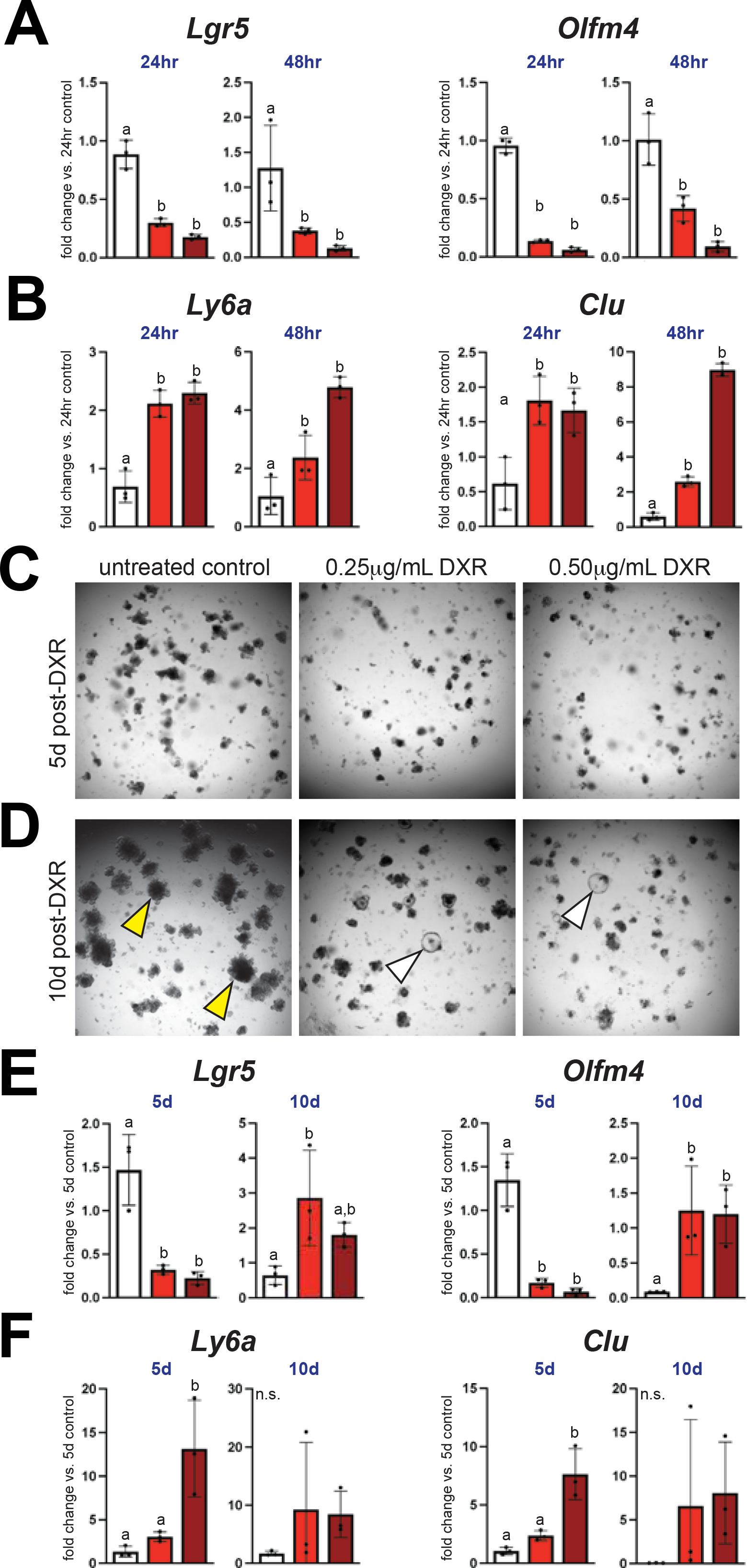
Doxorubicin downregulates aISC markers and induces a fISC response in intestinal organoids. (A) At 24hr and 48hr post-DXR, aISC markers *Lgr5* and *Olfm4* are significantly downregulated, while (B) fISC markers *Ly6a* and *Clu* are significantly upregulated. (C) At 5d post-injury, DXR treated organoids remain small and surrounded by debris. (D) By 10d post-injury, cystic organoids are visible in DXR treated wells (white arrowheads), while controls have extensive budding morphology consistent with differentiation (yellow arrowheads) (scale bar represents 100μm). (E) aISC markers remain significantly downregulated relative to untreated controls at 5d post-DXR, and are upregulated at 10d post-DXR, consistent with proliferative, cystic morphology. (F) While fISC markers remain elevated in organoids treated with 0.50μg/mL DXR 5d after injury, variable expression is observed at 10d post-DXR (n = 3 replicates per group, different letters indicate significant differences between sample groups at each timepoint, n.s. indicates not significant, p < 0.05).

To ask if organoids can return to an aISC-dominant homeostasis following DXR, we conducted RT-qPCR for aISC and fISC markers at later post-injury timepoints. Morphologically, DXR-treated organoids remained small and surrounded by apoptotic cells at 5d, suggesting ongoing IEC damage response (Fig. 2C). Accordingly, both *Lgr5* and *Olfm4* remained significantly downregulated in both DXR conditions relative to matched untreated controls, while *Ly6a* and *Clu* were significantly elevated (Fig 2E & F). At 10d post-DXR, aISC markers were significantly elevated in DXR-treated samples relative to controls (Fig 2E). While this result was initially unexpected, we observed that DXR-treated organoids at this timepoint exhibit cystic morphology, consistent with high WNT signaling (Gracz et al., 2015; Sato et al., 2011). Control organoids were, by contrast, large and budded, consistent with more mature IECs (Fig 2D). Interestingly, fISC markers were not significantly upregulated at 10d post-DXR due to high well-to-well variability (Fig 2F). Together, these data suggest that organoids at 5d post-DXR are still in a regenerative, fISC-associated state. By 10d post-DXR, morphology and gene expression suggest that organoids are undergoing aISC expansion, with some heterogeneous, residual fISC gene expression.

### Persistent aISCs are lost at DXR doses incompatible with organoid survival

To track aISC and fISC dynamics after DXR injury, we conducted a preliminary experiment using organoids from *Lgr5*^*EGFP-IRES-CreER*^ mice (Barker et al., 2007). Our original intent was to monitor aISC loss and re-emergence over time via confocal microscopy. However, optimization experiments using widefield microscopy yielded the unexpected result that *Lgr5*^*high*^ cells were never fully lost in organoids surviving 0.50μg/mL DXR, though the rate of their expansion slowed relative to controls (Fig S3A). While these results suggest that some aISCs persist through injury, *Lgr5*^*EGFP*^ expression is mosaic and accurately quantifying individual ISCs by microscopy is technically challenging and error prone. We sought to confirm our observation using an alternative approach that would facilitate quantitative monitoring of aISC loss by flow cytometry.

Previous *in vivo* studies from our lab and others have demonstrated that differential expression levels of a *Sox9*^*EGFP*^ transgene can identify aISCs (*Sox9*^*low*^; Sox9L), transit-amplifying (TA) progenitors (*Sox9*^*sublow*^; Sox9SL), secretory progenitors and mature enteroendocrine (EEC) and tuft cells (*Sox9*^*high*^; Sox9H), and all other mature IEC lineages (*Sox9*^*neg*^; Sox9N) (Formeister et al., 2009; Raab et al., 2020). While *Sox9*^*EGFP*^ organoids express heterogeneous EGFP levels, similar to *in vivo*, whether these levels correspond to the same cell populations has not been shown (Fig 3A). We isolated Sox9N, Sox9SL, Sox9L, and Sox9H populations by FACS and conducted RT-qPCR (Fig S4). As observed *in vivo*, we found that the Sox9L population was enriched for aISC marker *Lgr5*, Sox9H was enriched for EEC marker *Chgb*, and Sox9N was enriched for the mature absorptive enterocyte marker *Fabp1* (Fig 3B). Consistent with progenitor identity, Sox9SL organoid cells expressed intermediate levels of *Lgr5* and *Fabp1* (Fig 3B). These results demonstrate that the *Sox9*^*EGFP*^ allele can capture the aISC-to-mature IEC spectrum of differentiation in organoids.

**Figure 3:**
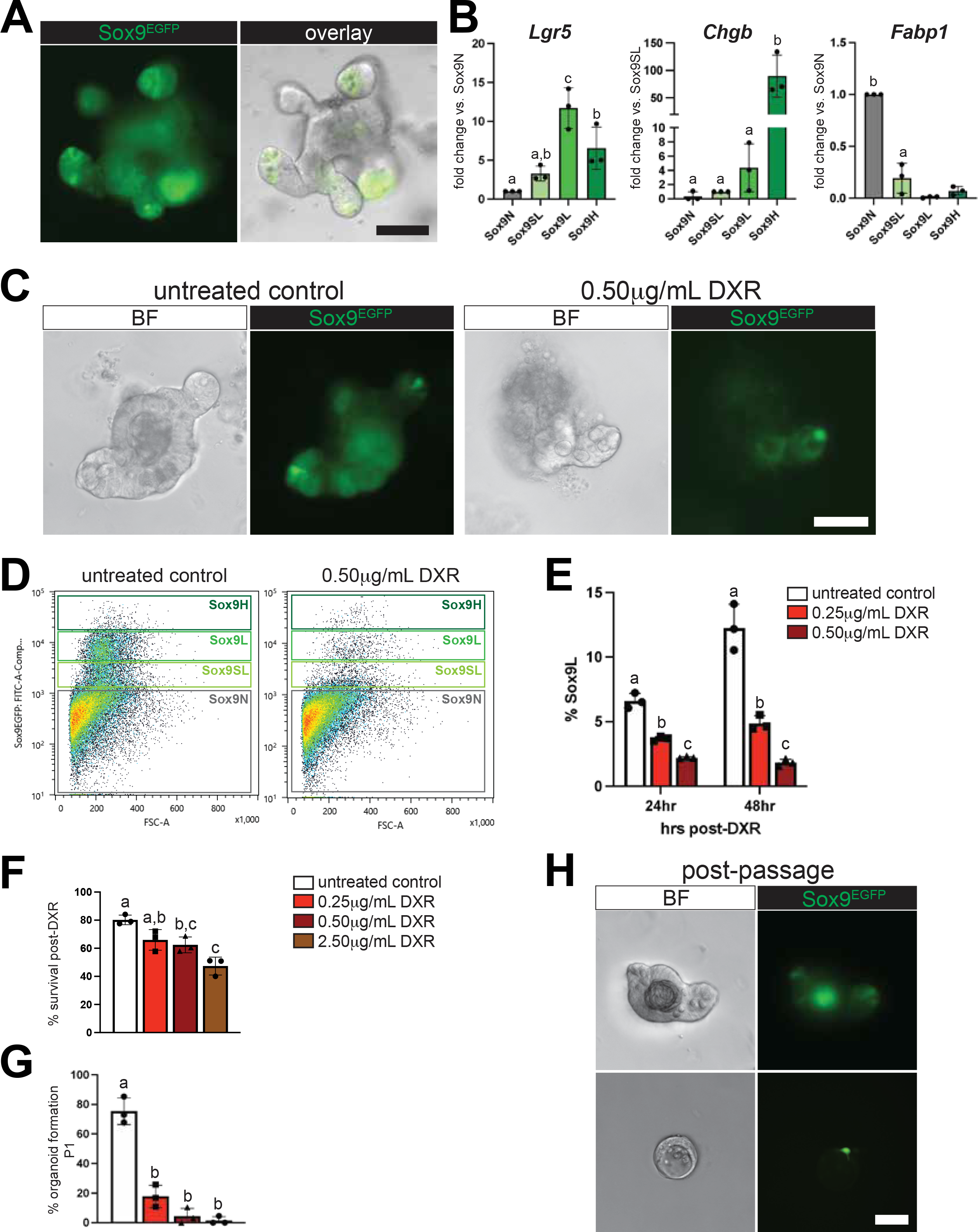
Residual aISCs persist through doxorubicin injury. (A) Sox9^EGFP^ is expressed at variable levels and enriched in crypt buds of organoids. (B) RT-qPCR validates that Sox9^EGFP^ populations in organoids are consistent with those observed *in vivo*, notably that Sox9L is significantly enriched for *Lgr5*. (C) DXR treatment results in a loss of EGFP+ cells and expected changes in organoid morphology. (D) Flow analysis validates loss of Sox9SL progenitors and Sox9L aISCs at 24hr post-DXR, which is incomplete and leaves residual aISCs. (E) DXR treatment results in dose-dependent aISC loss and stunts expansion of aISCs from 24hr to 48hr post-DXR. (F) P0 organoids exhibit dose-dependent decreases in survival at 24hr post-DXR and (G) dose-dependent loss of organoid forming capacity from single cells at P1. (H) Representative images of organoids, 5d post-passage as single cells from DXR-treated organoids at 24hr (n = 3 replicates per sample group, different letters indicate significant differences between sample groups at each timepoint, p < 0.05) (scale bar in 3H represents 50μm; all other scale bars represent 100μm).

Next, we treated *Sox9*^*EGFP*^ organoids with 0.25μg/mL and 0.50μg/mL DXR to monitor aISC loss. As expected, DXR treatment induced cell death and reduced the number of visible EGFP+ cells in organoids (Fig 3C). When we analyzed organoids by flow cytometry 24hr and 48hr after DXR, we observed a dose-dependent decrease in Sox9L aISCs relative to untreated controls (Fig 3D & E). However, even organoids treated with 0.50μg/mL DXR were composed of 1.8% ± 0.3% aISCs at 48hr post-DXR. Like preliminary observations made in Lgr5^EGFP^ organoids, % Sox9L aISCs expanded in untreated control cultures between 24hr and 48hr but remained unchanged between these timepoints following DXR (Fig 3E). Proliferative Sox9SL TAs were also sensitive to DXR treatment, and Sox9N cells increased by proportion of total viable IECs, reflecting a shift from ISC/TA populations to mature IECs (Fig S3B). To determine if DXR concentrations incompatible with organoid survival induce even greater aISC loss, we conducted an independent experiment treating organoids with 0.25, 0.50, 1.00, and 2.50 μg/mL DXR. By fluorescence microscopy, Sox9SL and Sox9L IECs were rare following 1.00μg/mL DXR and only Sox9H and Sox9N IECs were visible after 2.50μg/mL DXR (Fig S3C). As expected, Sox9N were resistant to DXR across all concentrations tested, consistent with post-mitotic status (Fig S3D). Sox9H exhibited minimal sensitivity to the highest DXR doses tested, which may reflect mature EEC or tuft cell identity, or known resistance to genotoxic injury via very low proliferation (Fig S3D) (Roche et al., 2015).

Next, we passaged organoids as single cells into WENR media at 24hr after DXR. We reasoned that the ability to form organoids after passage would provide functional evidence for aISC presence post-DXR injury. DXR induced expected levels of organoid loss at 24hr at P0 (Fig 3F). At 5d post-passaging, IECs from control organoids robustly generated new organoids, while IECs from DXR-treated organoids generated significantly fewer and exhibited impaired morphology (Fig 3G & H). While differences in P1 organoid formation between DXR-treated sample groups did not reach statistical significance, organoids were present in all three replicates from P0 organoids treated with 0.25μg/mL DXR but only one organoid from P0 organoids treated with 2.50μg/mL DXR was observed across all replicate wells (Fig. 3G). These data demonstrate that functional stemness is retained at lower doses of DXR, and almost completely lost at DXR doses incompatible with organoid survival. Together with quantification of aISCs in *Sox9*^*EGFP*^ organoids, they suggest that some aISCs survive initial DXR injury in a dose-dependent manner and that the presence of a persistent aISC pool correlates with organoid survival.

### Lgr5+ aISCs are required for organoid survival after DXR

To directly test if aISCs are required for organoid survival, we generated organoids from *Lgr5*^*2A-DTR/+*^ mice. This allele allows for ablation of *Lgr5+* aISCs by administration of diphtheria toxin (DT) and is expressed robustly in all aISCs (Tan et al., 2021). We first tested our ability to ablate *Lgr5*+ aISCs without compromising organoid survival. Organoids were grown for 72hr, to match the timepoint corresponding with 24hr post-injury in our DXR model, then “pulsed” with 100ng/mL DT for 1, 2, or 3hr. *Lgr5* was decreased ∼5-fold by RT-qPCR at 24hr post-DT regardless of the length of DT treatment (Fig. S5A). We next pulsed organoids with DT for 3hr and quantified survival at 5d post-DT. DT administration induced morphological changes in organoids consistent with cell death, and organoids were surrounded by dead cells and debris at 48hr post-DT and onward (Fig S5B). However, DT-mediated ablation of *Lgr5*+ cells did not significantly decrease survival at 24hr or 120hr (Fig SC). Together, these data validate *Lgr5*^*2A-DTR*^ organoids as a tool for ablating aISCs without compromising organoid survival.

Next, we designed an experiment to ablate residual aISCs in DXR-treated organoids at 24hr post-injury (Fig. 4A). We selected 0.25 μg/mL DXR, reasoning organoids suffering less genotoxic injury should be more resistant to aISC ablation. *Lgr5*^*2A-DTR/+*^ organoids treated with DXR only were lost at the same rate as in previous experiments and demonstrated similar morphological signs of injury (Fig. 4B & C). While DT-treated organoids exhibited a small drop in survival at early timepoints, survival we not significantly different relative to controls at 5d post-DT (Fig. 4C). In contrast, organoids treated sequentially with DXR and DT exhibited extensive damage and decreased survival, with no surviving organoids observed at 5d post-DT (6d post-DXR) (Fig. 4B & C). RT-qPCR on organoids 24hr after DT treatment (48hr post-DXR) demonstrated significantly decreased *Lgr5* and *Olfm4* in all treatment groups with no differences between DXR, DT, or DXR+DT, while *Ascl2* was significantly downregulated following DXR+DT vs. DT alone (Fig. 4D & E). All treatment conditions also upregulated fISC markers *Ly6a* and *Clu*, though *Clu* was not as elevated in organoids treated only with DT relative to those treated with DXR and DXR+DT (Fig. 4F). Collectively, DT ablation experiments demonstrate that residual aISCs surviving initial genotoxic injury are required for subsequent organoid survival, even at lower doses of DXR. Further, our data show that upregulation of “canonical” fISC genes is an insufficient readout of fISC function in organoid models of IEC injury and suggest that regeneration independent of the aISC pool may rely on non-epithelial signaling or exogenous factors not present in standard organoid media.

**Figure 4:**
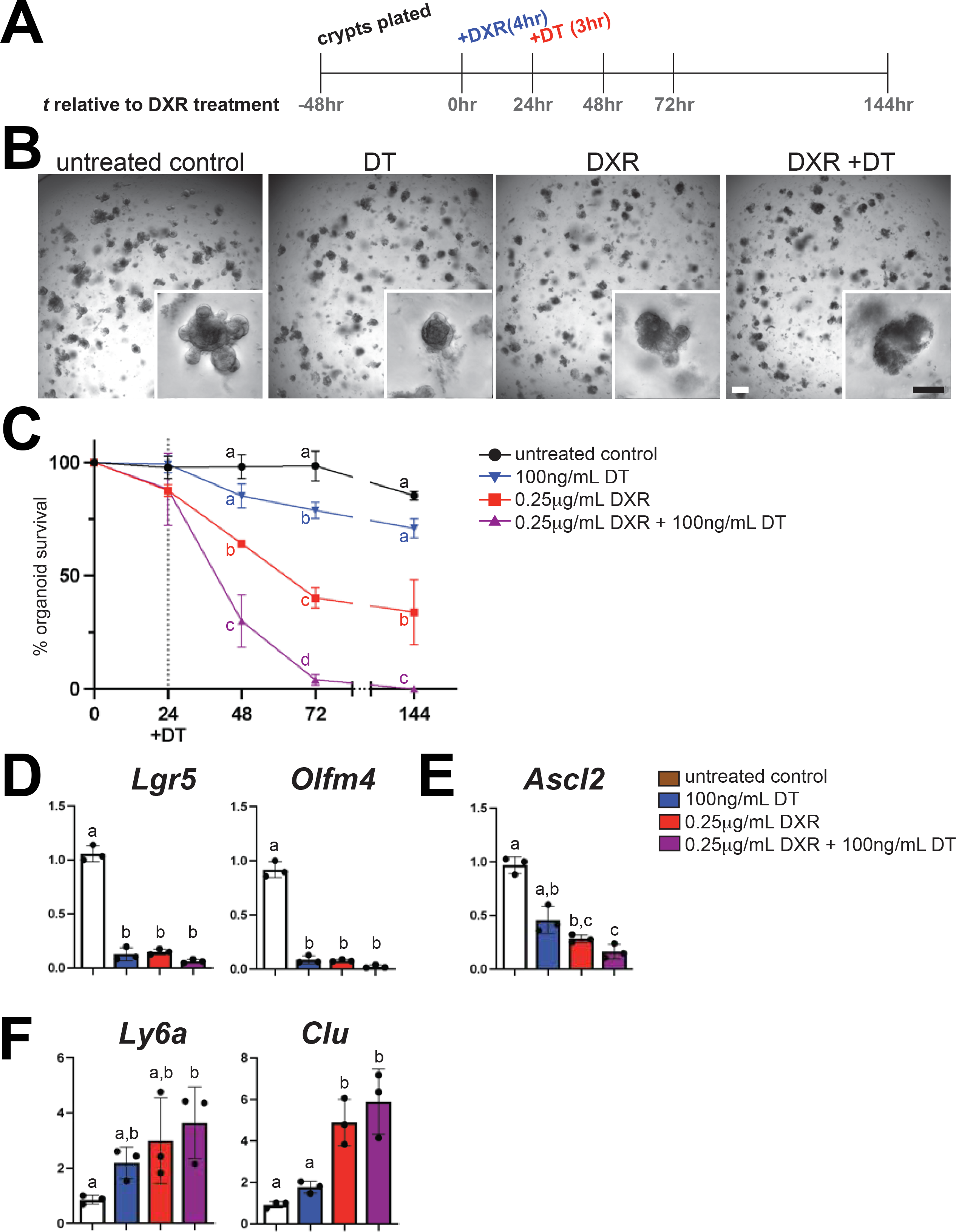
Surviving aISCs are required for organoid regeneration after doxorubicin injury. (A) DT treatment 24hr after DXR aims to ablate residual aISCs in *Lgr5*^*2A-DTR*^ organoids. (B) Post-DXR ablation of *Lgr5*+ ISCs results in extensive morphological damage (white scale bar represents 250μm; black scale bar represents 100μm) (C) Loss of *Lgr5*+ ISCs eliminates the ability of organoids to survive 0.25μg/mL DXR (D) aISC markers *Lgr5* and *Olfm4* are downregulated following either DXR or DT treatment alone, but undetectable in organoids treated with DT 24hr after DXR injury. (E) *Ascl2* is downregulated in all injury conditions, but significantly reduced in DXR+DT relative to untreated organoids and organoids treated with DT alone. (F) fISC markers are upregulated in response to DXR but exhibit more modest and gene-dependent upregulation in response to DT alone (n = 3 replicates per group, different letters indicate significant differences, p < 0.05).

### Conclusions

Our results highlight significant differences in aISC/fISC responses to genotoxic injury between organoids and *in vivo* models and suggest that careful consideration of these differences is critical for interpretation of certain *in vitro* assays. Studies designed to screen aISC/fISC-related functions, such as the ability of chemotherapeutic drugs to inhibit tumor growth while sparing normal epithelial cells, may need to consider absent or impaired fISC function in intestinal organoids. Recent reports demonstrated that co-culturing patient-derived colon cancer organoids with cancer-associated fibroblasts resulted in decreased sensitivity to genotoxic injury, highlighting non-epithelial contributions to organoid damage response (Ramos Zapatero et al., 2023). Our findings that *Lgr5*+ aISCs are required for intestinal organoid regeneration after DXR injury are also consistent with growing evidence that epithelial-stromal interactions are critical for the fISC response. TGF-β derived from the stromal compartment was recently shown to induce the fISC state *in vitro* and contribute to post-injury intestinal regeneration *in vivo* (Chen et al., 2023). Though outside of the scope of the current study, modified organoid conditions modeling the damaged ISC niche, might better model regeneration *in vitro*. Clarification of the molecular and genetic requirements for fISC function in organoids is necessary to expand the utility of these models. The optimized model described here will facilitate further studies into aISC/fISC dynamics *in vitro*.

## Acknowledgements

The authors thank members of the Gracz lab for helpful conversations and critical review of the manuscript. This work was funded by NIH R35GM142503 (Gracz) and R35GM142503-01S1 (Gracz). This work was supported in part by the Emory Mouse Transgenic and Gene Targeting Core (TMF), for which additional support was provided by the National Center for Advancing Translational Sciences of the NIH (UL1TR000454).

