## Supplemental Materials for "*Lgr5*+ intestinal stem cells are required for organoid survival after genotoxic injury"

**Lee, et al.**

### SUPPLEMENTAL FIGURE LEGENDS

**Figure S1:** ***Doxorubicin treatment and dose response in intestinal organoids.*** (A) For DXR treatment, crypts are allowed to establish as organoids in ENR media for 48hr prior to a 4hr pulse of DXR. In initial dose curve studies, organoid survival was quantified daily to 6d post-DXR. (B) DXR dose curve identifies 0.25μg/mL and 0.50μg/mL as optimal doses compatible with long-term survival. (C) Statistical comparisons of organoid survival at each post-injury timepoint demonstrate dose-dependent effects of DXR treatment (n = 3 replicates per group, different letters indicate significant differences between sample groups at each timepoint, $p < 0.05$).

**Figure S2:** ***Ascl2 response to doxorubicin treatment.*** *Ascl2* is significantly downregulated by DXR at 24hr, but decreased expression in untreated organoids eliminates significant differences at 48hrs. *Ascl2* expression is not significantly different relative to untreated controls at 5d post-DXR, but is significantly upregulated at 10d post-DXR, resembling *Lgr5* and *Olfm4* expression (Fig 2E) (n = 3 replicates per group, different letters indicate significant differences, n.s. indicates not significant, $p < 0.05$).

**Figure S3:** ***aISCs exhibit a dose-dependent response to doxorubicin in organoids.*** (A) Quantification of $Lgr5^{high}$ cells in individual $Lgr5^{EGFP\text{-}IRES\text{-}CreER}$ organoids suggests that the aISC pool is never fully lost in organoids that survive DXR treatment, but rather that DXR inhibits aISC expansion through 5d post-injury (asterisks indicate significant differences between untreated controls and DXR treated organoids at each timepoint, $p < 0.05$). (B) Flow cytometry data for all Sox9EGFP populations corresponding to Fig 3E. (C) High doses of DXR result in organoids consisting mainly of Sox9H and Sox9N IECs, with Sox9SL and Sox9L populations very rare following 2.50μg/mL DXR (representative images show 24hr post-DXR, scale bar represents 100μm). (D) DXR treatment results in dose-dependent loss of Sox9SL TAs and Sox9L aISCs at 24hr post-injury by flow cytometry (n = 3 replicates per group, different letters indicate significant differences, $p < 0.05$).

**Figure S4:** ***Gating strategy for flow/FACS of Sox9EGFP populations.*** Representative flow cytometry plots are shown for untreated control and 0.25μg/mL DXR organoids at 24hr post-DXR.

**Figure S5:** ***Validation of $Lgr5^{2A\text{-}DTR}$ organoids for aISC ablation.*** (A) 1, 2, and 3hr "pulse" of DT significantly downregulates *Lgr5* expression in $Lgr5^{2A\text{-}DTR}$ mice at 24hr (n = 3 replicates per group, different letters indicate significant differences, $p < 0.05$). (B) DT treatment impacts organoid morphology, resulting in cell death and accumulation of debris, but typical budding morphology returns by 5d post-DT (white scale bar represents 250μm; black scale bar represents 100μm). (C) DT treatment does not impact $Lgr5^{2A\text{-}DTR}$ organoid survival (n = 3 replicates per group, n.s. indicates not significant, $p < 0.05$).

### METHODS

#### *Mice*

All experiments were carried out using mice between 8 and 24 weeks of age, maintained on C57Bl/6 background. Sox9$^{EGFP}$ (Gong et al., 2003), Lgr5$^{EGFP\text{-}IRES\text{-}CreER}$ (Jackson Labs, strain #008875) (Barker et al., 2007), and Lgr5$^{2A\text{-}DTR}$ (Tan et al., 2021) mice have been previously described. All transgenic/mutant alleles were maintained at heterozygosity. Animals received PicoLab Rodent Diet 20 (LabDiet, 5053) and water *ad libitum*. The Emory University Institutional Animal Care and Use Committee reviewed and approved all animal protocols.

#### *Crypt isolation and organoid culture*

Crypts were isolated for organoid culture as previously described (Zwarycz et al., 2018). Briefly, intestines were dissected out, the first 6cm discarded, and the remaining proximal half designated as jejunum and taken for organoid isolation. Jejunal segments were opened longitudinally and rinsed briefly in a 50mL conical containing 10mL sterile Dulbecco's Phosphate-Buffered Saline (DPBS) (Gibco, 14190250). Tissue was transferred to a new 50mL conical containing 3mM EDTA (Corning, 46-034-CI) in 10mL sterile DPBS and incubated at 4C on a rocking platform set to 40RPM for 15min. Tissue was retrieved and villi removed by gentle "brushing" with a P200 pipette tip on a glass plate. Intestinal tissue was then rinsed briefly in a petri dish containing sterile DPBS and cut into pieces ~0.5cm in length before being transferred to a new 50mL conical containing 3mM EDTA in 10mL sterile DPBS and incubated at 4C on a rocking platform for 35min. Jejunal pieces were transferred to a 50mL conical containing 10mL sterile DPBS and shaken for 3-4min to release crypts, confirming expected crypt density and morphology by light microscopy at 1min intervals to determine when to end the dissociation protocol. 10mL sterile DPBS was added to isolated crypts, which were then filtered through a 70 or 100µm cell strainer, pelleted at 600g for 5min at RT, and resuspended in 200-500uL Advanced DMEM/F12 (Gibco, 12634010).

Crypt density per 10uL media was examined qualitatively by light microscopy and crypt density estimated, with the goal of determining volume required to achieve between 50-100 crypts per 10µL Matrigel in culture. Crypts were resuspended in 75% phenol red-free, growth factor-reduced Matrigel (Corning, 356231) and plated as 10µL droplets in 96 well plates or 40µL droplets in 48 well plates. Matrigel was allowed to polymerize at RT for 5min, then at 37C for 15min. Media was overlaid at 100µL per well in 96 well plate and 200µL per well in 48 well plates: Advanced DMEM/F12, 1X N2 (Thermo Fisher, 17502048), 1X B27 w/o vitamin A (Thermo Fisher, 12587010), 1X HEPES (Gibco, 15630080), 1X Penicillin/Streptomycin (Sigma-Aldrich, P4333-100ML), 1X Glutamax (Gibco, 35050061), 10% RSPO1-CM (made using RSPO1 transfected HEK293T cells following manufacturer protocol: Sigma Aldrich, SCC111), 50ng/mL recombinant murine EGF (Gibco, PMG8041), and 100ng/mL recombinant human NOGGIN (PeproTech, 120-10C-20UG). 500µg/mL Primocin (Invivogen, ant-pm-1) and 10mM Y27632 (Selleck Chemicals, S1049) was added to overlay media for the first 48hr after plating crypts and then excluded from all other media changes. ENR media was replaced every 48 hours.

#### *Doxorubicin and diphtheria toxin treatment*

Intestinal organoids were treated with doxorubicin (Selleck Chemicals, S1208) 48 hours after plating. Doxorubicin was added and mixed twice in ENR media by gentle pipetting. After incubating for four hours at 37°C, the treated media was removed and discarded, and organoid wells were washed with 150µL of warmed DPBS for 10 minutes at 37°C, twice. After washing, each well received 100µL fresh ENR media.

Organoids were treated with 100ng/mL diphtheria toxin (Cayman Chemical, 19657-1) 72 hours after plating. Diphtheria toxin was added into ENR overlay media and mixed by gently pipetting twice. After incubating for three hours at 37°C, DT media was removed and discarded and DT-treated wells received two 10min washes with 150µL of DPBS at 37°C. 100µL of new ENR media was overlaid into each well following DPBS washes.

For pilot studies using *Lgr5$^{EGFP\text{-}IRES\text{-}CreER}$* organoids, DXR treatment protocols and timelines were followed as described above. Organoids were imaged on an Olympus IX-81 inverted epifluorescence microscope with cellSens software (Olympus). Individual organoids were identified prior to DXR treatment and stage positions programmed in order to quantify cells in the same organoid day-to-day. To account for transgene mosaicism, only EGFP+ organoids were selected for ongoing quantification. *Lgr5$^{high}$* cells were quantified by manually examining each organoid through its entire *z* plane and counting clear, wedge-shaped EGFP$^{high}$ cells that were distinct from surrounding background fluorescence.

***Passaging for single cell organoid formation assay***
Organoid passaging was performed 24h after DXR treatment (72h after plating). Each well of a 96 well plate was resuspended in 200 µl of TrypLE and incubated 3 minutes at 37°C in water bath. TrypLE was quenched with 1mL cold 1% BSA in DPBS and epithelial fragments mechanically dissociated by pipetting. After centrifugation at 600g for 5 minutes at 4°C, the pellet was washed with 1mL cold 1% BSA in DPBS. After centrifugation, the pellet was resuspended, plated in 75% Matrigel suspension in a 48 well plate, and allowed to polymerize at 37C for 15min. 200µL WENR media per well was overlaid: 50% Advanced DMEM/F12, 40% WNT3A-CM (made using L-WNT3A cells following manufacturer protocol: ATCC CRL-2647), 10% RSPO1-CM, 1X N2, 1X B27 w/o vitamin A, 1X HEPES, 1X Penicillin/Streptomycin, 1X Glutamax, 50ng/mL recombinant murine EGF, and 100ng/mL recombinant human NOGGIN. 500µg/mL Primocin and 10mM Y27632 were added to overlay media for the first 48hr after passaging and then excluded from all other media changes. Culture was maintained for 5d before counting grown organoids by light microscopy.

***Flow cytometry and FACS***
Organoids were dissociated for flow analysis and FACS by removing overlay media and pooling 6-8 wells per sample from a 48 well plate into a single 5mL conical with 1% BSA in DPBS. Pooled organoids were pelleted at 600g for 5min, then resuspended in 1mL TrypLE and incubated at 37C for 5min. TrypLE was quenched by adding 2mL Advanced DMEM/F12 and organoids pipetted 25X with a P1000 to aid in mechanical dissociation. Dissociated organoid cells were resuspended in 1mL Advanced DMEM/F12 with 5uL each 7-AAD (Biolegend, 420404) and Annexin V-APC (Biolegend, 640941) for live/dead discrimination. Flow analysis and FACS were carried out on a Sony SH800 (Sony Biotechnology). To quantify distribution of *Sox9$^{EGFP}$* populations, cells were gated for doublet-discrimination and to exclude dead cells. EGFP populations were calculated as percent of total viable singlets (see Fig. S4 for gating). For RNA isolation, cells were sorted

directly into 500uL Lysis Buffer (Ambion RNAqueous Micro Kit, AM1931) and stored at -70C until further processing (see ***RNA isolation and RT-qPCR***).

***RNA isolation and RT-qPCR***

Organoids were lysed at various timepoints after treatment using 200µL of Lysis Buffer from the RNAqueous Micro Total RNA Isolation Kit (Ambion, AM1931). Organoid lysis was completed by gently pipetting lysed wells 3-5 times in Lysis Solution, prior to storage overnight or longer at -70C. RNA was purified using the Ambion RNAqueous Micro kit, according to manufacturer instructions. All samples were treated with DNase (Ambion) for 30min at 37C and DNase inactivated following manufacturer protocol. Total RNA was quantified using a Qubit v3 Fluorometer (Thermo Fisher, Q33216) and the Qubit High Sensitivity RNA Quantification Assay (Thermo Fisher, Q32855). 50-100ng RNA was subjected to reverse transcription using the iScript cDNA Synthesis Kit (Bio-Rad 1708891) and diluted 1:5 in molecular grade water (Corning, 46-000-CI)). RT-qPCR was carried out in technical triplicate using Taqman assays and SsoAdvanced Universal Probes Supermix (BioRad, 1725284), following manufacturer protocols and using 1uL diluted cDNA input per reaction. Reactions were run on a QuantStudio 3 Real Time PCR instrument (Thermo Fisher) and analyzed using the $\Delta\Delta C_T$ method (Pfaffl, 2001). *Actb* was selected as the internal housekeeping gene. Taqman assay IDs used in this manuscript are: *Actb* (Mm02619580_g1), *Ascl2* (Mm01962673_s1), *Clu* (Mm01197002_m1), *Lgr5* (Mm00438890_m1), *Ly6a* (Mm00725565_m1), and *Olfm4* (Mm01320260_m1).

**Barker, N., van Es, J. H., Kuipers, J., Kujala, P., van den Born, M., Cozijnsen, M., Haegebarth, A., Korving, J., Begthel, H., Peters, P. J., et al.** (2007). Identification of stem cells in small intestine and colon by marker gene Lgr5. *Nature* **449**, 1003-1007.

**Gong, S., Zheng, C., Doughty, M. L., Losos, K., Didkovsky, N., Schambra, U. B., Nowak, N. J., Joyner, A., Leblanc, G., Hatten, M. E., et al.** (2003). A gene expression atlas of the central nervous system based on bacterial artificial chromosomes. *Nature* **425**, 917-925.

**Pfaffl, M. W.** (2001). A new mathematical model for relative quantification in real-time RT-PCR. *Nucleic Acids Res* **29**, e45.

**Tan, S. H., Phuah, P., Tan, L. T., Yada, S., Goh, J., Tomaz, L. B., Chua, M., Wong, E., Lee, B. and Barker, N.** (2021). A constant pool of Lgr5(+) intestinal stem cells is required for intestinal homeostasis. *Cell Rep* **34**, 108633.

**Zwarycz, B., Gracz, A. D. and Magness, S. T.** (2018). Organoid Cultures for Assessing Intestinal Epithelial Differentiation and Function in Response to Type-2 Inflammation. *Methods in molecular biology* **1799**, 397-417.
