## Supplementary figures and images for "*Lgr5*+ intestinal stem cells are required for organoid survival after genotoxic injury"

### Figure S1

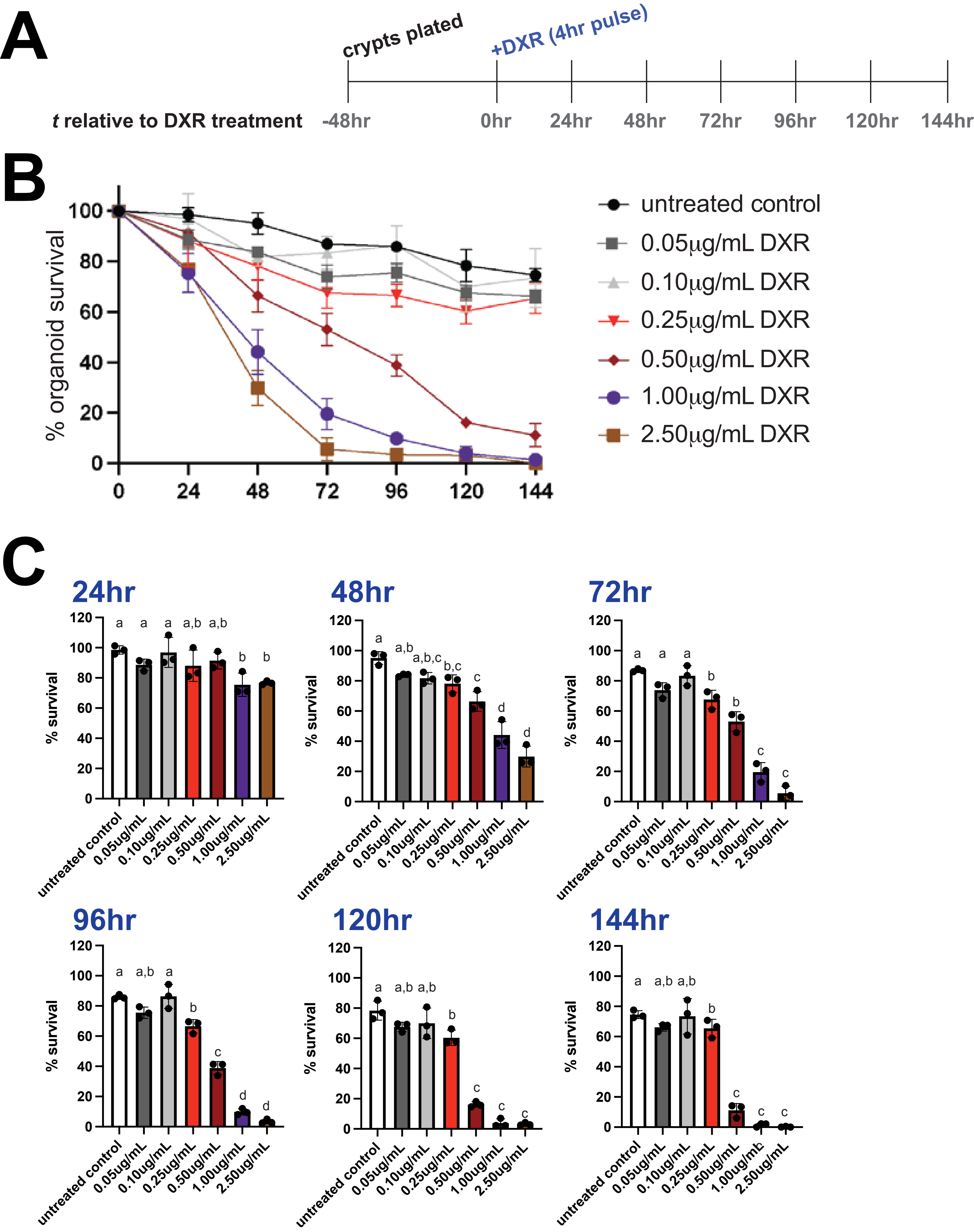

### Figure S2

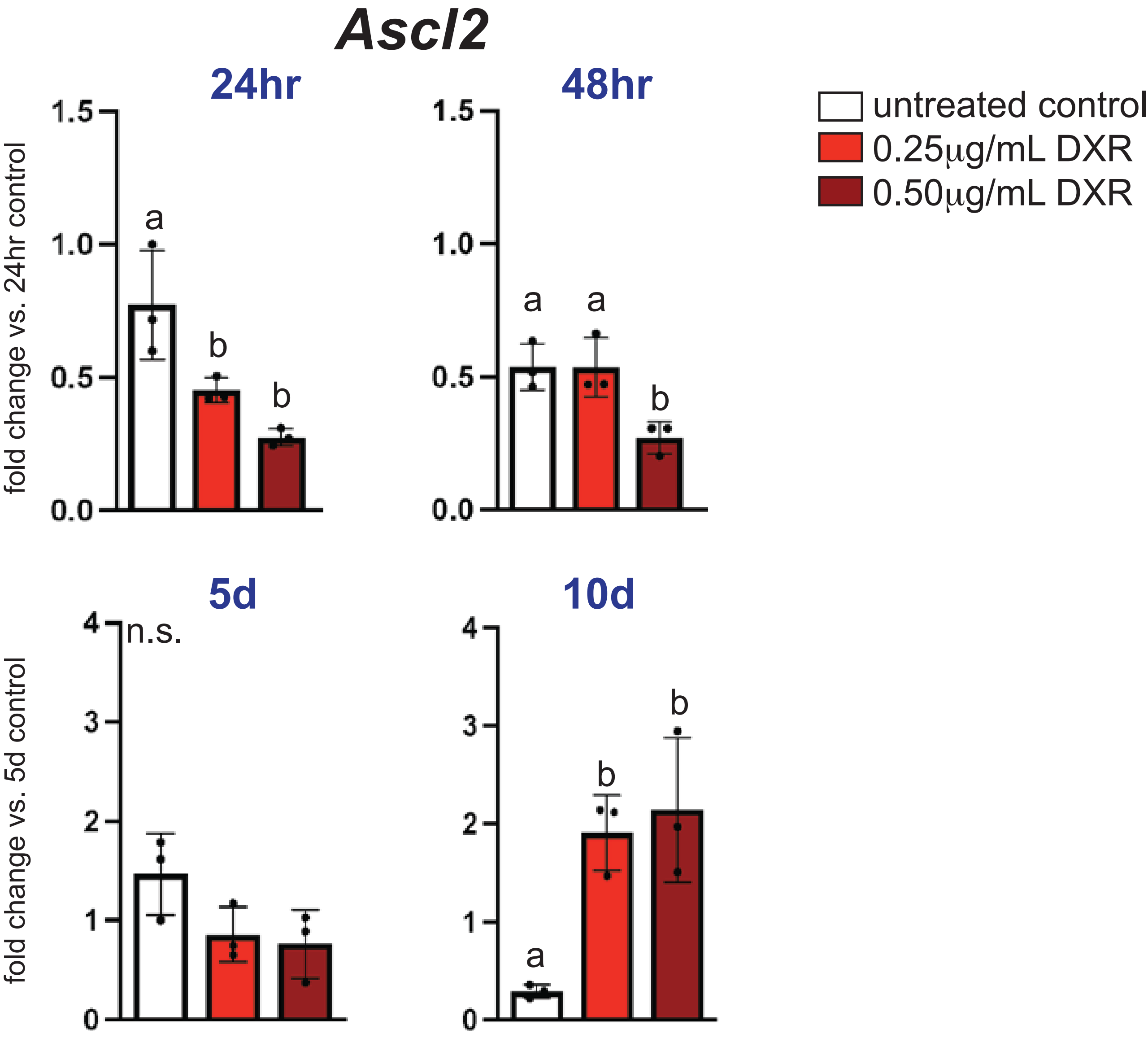

### Figure S3

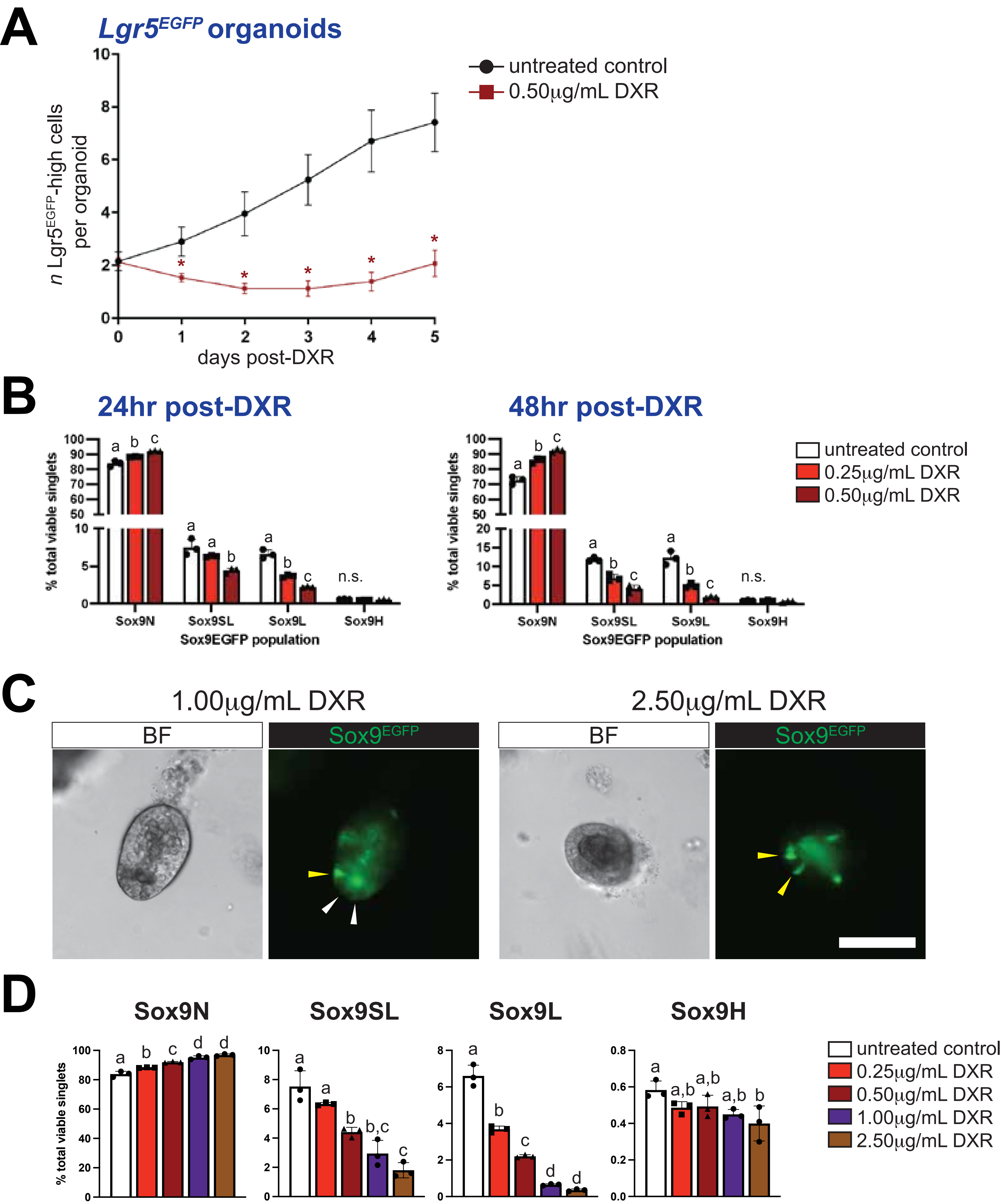

### Figure S4

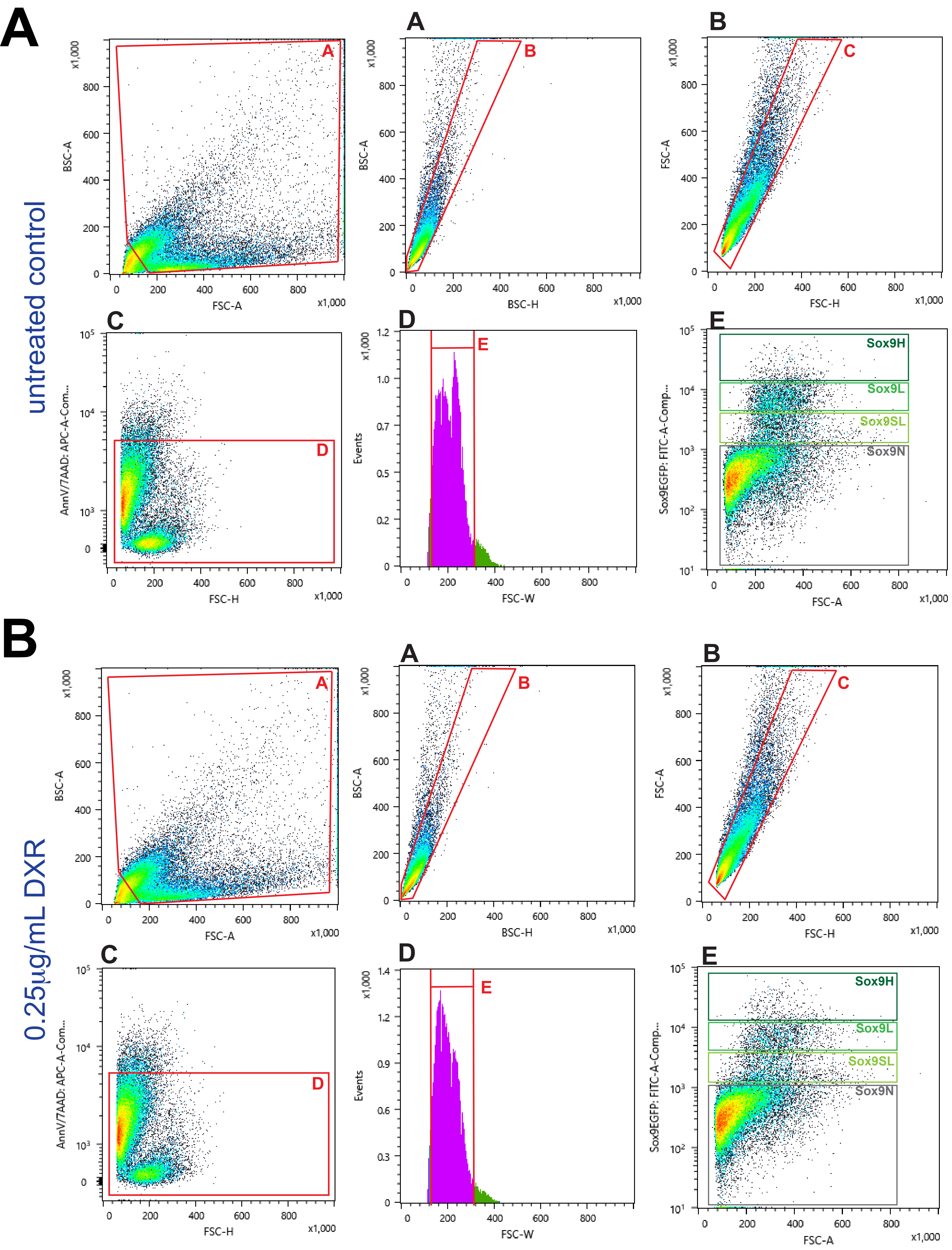

### Figure S5

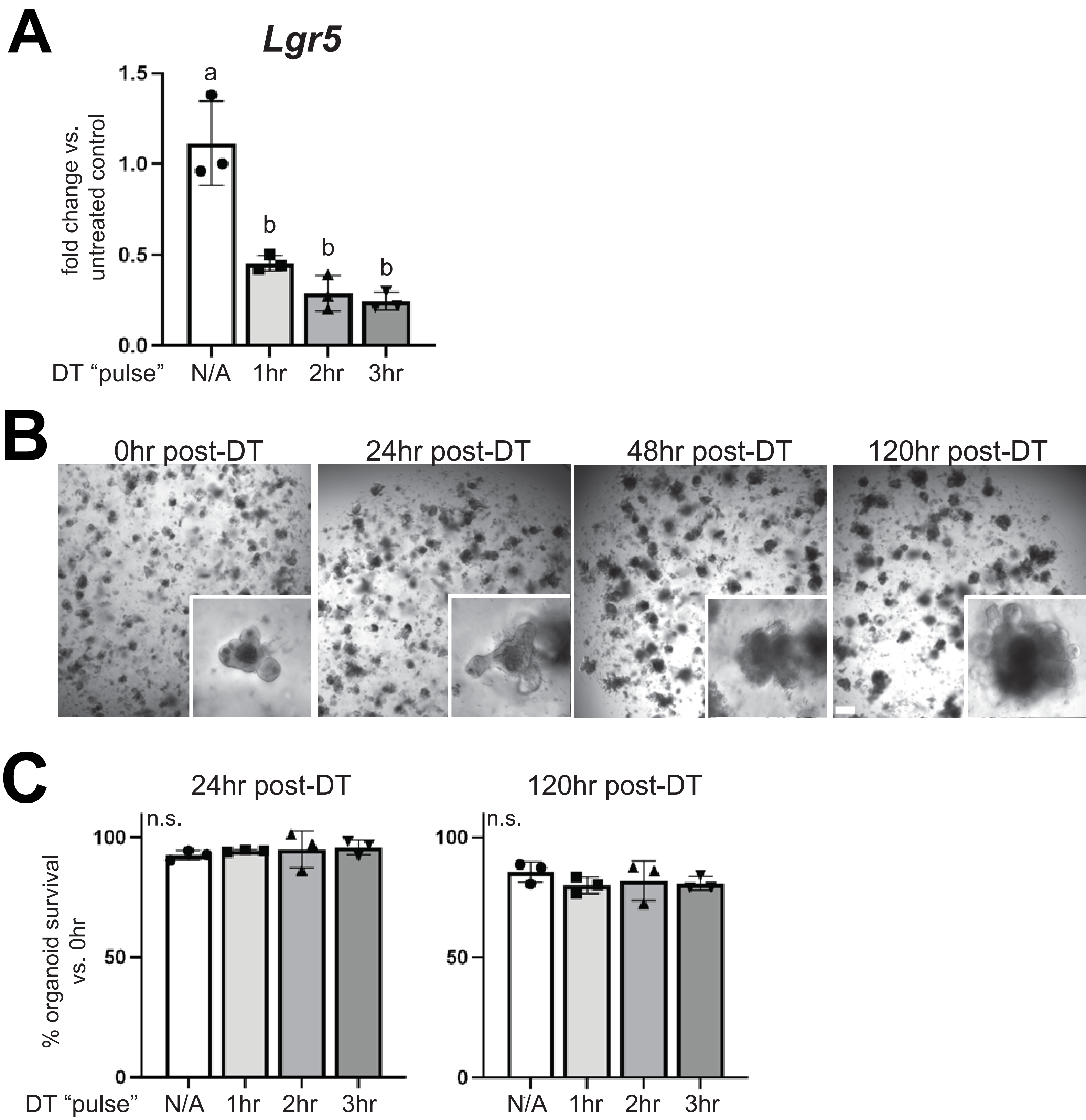
